# Laboratory-Based Surveillance of Animal Rabies in the Bono East Region of Ghana: A Six-Year Retrospective Study (2020 – 2025)

**DOI:** 10.64898/2026.07.12.738119

**Authors:** Prince Kyere Dwaah, Sylvia Afriyie Squire, Helen Djang-Fordjour, Issah Bagulo, Anthony Osei-Tutu, Samuel Opokuware, Emmanuel Opoku, Roger Twum Gyarko, David Kando, Abraham Num, Princess Amissah, Clement Amoah Newton Ntow, Nana Yaa Awua-Boateng

## Abstract

**Background:** Dog-mediated rabies remains a major neglected tropical disease and an important cause of preventable human mortality in sub-Saharan Africa. Despite ongoing control efforts, limited laboratory-confirmed surveillance data hinder accurate assessment of the disease burden and the development of evidence-based interventions in Ghana. This study investigated the epidemiological characteristics of laboratory-confirmed rabies cases among suspected animal submissions and evaluated the contribution of routine diagnostic surveillance to rabies control.

**Methodology/Principal Findings:** A retrospective analysis was conducted using laboratory surveillance records of suspected rabies cases submitted to a regional veterinary laboratory in Ghana between 2020 and 2025. Brain tissue samples were examined using Seller’s staining, the direct fluorescent antibody test (FAT), and polymerase chain reaction (PCR). Epidemiological information, including animal species, vaccination history, ownership status, bite history, and diagnostic outcomes, was extracted and analysed descriptively. Of the 51 suspected cases examined, 46 (90.2%) were laboratory confirmed as rabies. Domestic dogs accounted for 96.1% of submissions, while cats represented 3.9%. Most suspected animals were either unvaccinated (64.7%) or had unknown vaccination status (33.3%), and 90.2% had reportedly bitten humans before submission. Confirmed cases were detected throughout the study period, indicating sustained viral circulation and continued zoonotic risk within the study area.

**Conclusions/Significance:** The consistently high proportion of laboratory-confirmed rabies cases demonstrates persistent transmission of rabies virus among domestic animals and highlights substantial risks to human health. The integration of conventional and advanced laboratory diagnostic methods strengthened surveillance and improved case confirmation. These findings underscore the urgent need to expand dog vaccination coverage, strengthen laboratory-based surveillance, improve bite reporting systems, and enhance collaboration between veterinary and public health sectors. Reinforcing integrated One Health strategies will be essential to accelerate progress towards the global goal of eliminating dog-mediated human rabies by 2030.

**Author Summary:** Rabies remains one of the world’s oldest and most devastating zoonotic diseases, which is fatal yet preventable, causing thousands of human deaths each year, particularly in Africa and Asia. In Ghana, domestic dogs are the principal reservoir of the rabies virus, but information on the epidemiology of animal rabies from routine laboratory surveillance is limited in many regions. We reviewed six years (2020 – 2025) of laboratory records from the Regional Veterinary Laboratory in Techiman, Bono East Region, to describe the epidemiological characteristics of suspected animal rabies cases and laboratory-confirmed infections. Of the 51 brain samples submitted for diagnosis, 46 (90.2%) were confirmed positive for rabies. Domestic dogs accounted for almost all submissions, and most confirmed cases involved animals that were either unvaccinated or had an unknown vaccination history. Furthermore, most suspected animals had reportedly bitten humans before laboratory investigation, highlighting the continuing public health risk posed by canine rabies. The study also documents how regional diagnostic capacity was strengthened through collaboration with national reference laboratories, enabling the routine use of internationally recommended diagnostic methods such as the fluorescent antibody test and polymerase chain reaction. These findings provide valuable evidence to support enhanced rabies surveillance, increased mass dog vaccination, improved public awareness, and strengthened One Health collaboration. Such measures are essential for reducing human rabies deaths and supporting Ghana’s contribution to the global goal of eliminating dog-mediated human rabies by 2030.

## Introduction

Rabies is one of the oldest and most devastating zoonotic diseases, causing acute and progressive encephalitis that is almost invariably fatal once clinical signs develop. The disease is caused by viruses belonging to the genus *Lyssavirus* within the family *Rhabdoviridae* and is transmitted primarily through the saliva of infected mammals, most commonly via bites (Hampson et al., 2022; World Health Organization [WHO], 2023). Although rabies is entirely preventable through timely post-exposure prophylaxis (PEP) and effective vaccination of animal reservoirs, it continues to impose a substantial public health burden in many low- and middle-income countries (Diallo et al., 2019; Amicizia et al., 2024; Fotedar & Ravish, 2025; Ritchie et al., 2025; Fotedar & Shankariah, 2026; Pardeshi et al., 2026). According to Hampson et al. (2022), WHO (2023) and FAO et al. (2024), each year, an estimated 59,000 people die from dog-mediated rabies worldwide, with more than 95% of these deaths occurring in Africa and Asia, where access to preventive interventions and diagnostic services remains limited. Beyond its impact on human health, rabies causes considerable economic losses through livestock mortality, costs associated with PEP, animal vaccination programmes, and reduced productivity, disproportionately affecting vulnerable populations that depend on livestock and companion animals for their livelihoods (Undurraga et al., 2023).

Domestic dogs are responsible for approximately 99% of all human rabies infections globally and constitute the principal reservoir for virus transmission in endemic settings (WHO, 2023; Wallace et al., 2022). Sustained transmission within dog populations is largely driven by inadequate vaccination coverage, unrestricted movement of free-roaming dogs, poor ownership practices, and limited implementation of coordinated rabies control programmes. According to Diallo et al. (2019), Amicizia et al. (2024), Fotedar & Ravish (2025), Ritchie et al. (2025), Fotedar & Shankariah (2026), and Pardeshi et al. (2026), these challenges are further compounded by weak surveillance systems, insufficient laboratory diagnostic capacity, and low public awareness, resulting in substantial underreporting of both animal and human cases. Consequently, the reported burden of rabies in many endemic countries is believed to represent only a fraction of the true incidence of the disease.

Across sub-Saharan Africa, rabies remains endemic despite the availability of safe and effective vaccines for both humans and animals. The region continues to experience thousands of preventable human deaths annually, with children under 15 years of age accounting for a disproportionately large share of fatalities because of their frequent interactions with dogs and delayed access to post-exposure treatment (WHO, 2023). Several studies have demonstrated that sustained mass vaccination of at least 70% of the dog population is sufficient to interrupt rabies transmission and dramatically reduce human exposure (Lechenne et al., 2021; Sambo et al., 2022). However, achieving and maintaining this vaccination threshold remains difficult in many African countries because of financial constraints, inadequate veterinary infrastructure, competing public health priorities, and limited community participation in vaccination campaigns (Cleaveland et al., 2021; Undurraga et al., 2023).

The global commitment to eliminate dog-mediated human rabies by 2030, commonly referred to as the "Zero by 30" initiative, represents a collaborative effort led by the WHO, the Food and Agriculture Organization of the United Nations (FAO), and the World Organisation for Animal Health (WOAH). This strategy promotes integrated interventions that combine mass dog vaccination, improved surveillance, timely access to PEP, strengthened laboratory diagnosis, and enhanced collaboration between the human and animal health sectors (WHO, 2023; FAO et al., 2024). Central to this strategy is the One Health concept, which recognises that the health of humans, animals, and the environment is interconnected and that sustainable rabies control requires coordinated action across multiple disciplines (Destoumieux-Garzón et al., 2022). Countries that have successfully reduced or eliminated canine rabies have consistently demonstrated the effectiveness of integrated One Health approaches supported by strong surveillance systems and sustained political commitment (Lechenne et al., 2021).

Reliable surveillance forms the cornerstone of effective rabies prevention and control. Laboratory confirmation not only improves the accuracy of disease reporting but also supports outbreak investigations, guides public health interventions, and informs vaccination strategies. Unfortunately, passive surveillance systems remain the predominant approach in many endemic countries, where only animals exhibiting obvious clinical signs or involved in biting incidents are submitted for laboratory investigation. This reliance on passive reporting frequently results in delayed detection, geographical bias, and substantial underestimation of disease occurrence (Coetzer et al., 2019; Taylor et al., 2022; Keshavamurthy et al., 2024; Kahariri et al., 2025). Strengthening laboratory-based surveillance is therefore essential for generating accurate epidemiological data needed to guide national rabies control programmes.

The laboratory diagnosis of rabies has advanced considerably over recent decades. The direct fluorescent antibody test (FAT) remains the internationally recommended reference method for detecting rabies virus antigen in brain tissue because of its high sensitivity and specificity (Tenzin et al., 2020; Cappelari et al., 2022; Claassen et al., 2023; WOAH, 2023; Todoroki et al., 2025). Molecular techniques such as reverse transcription polymerase chain reaction (RT-PCR) have further improved diagnostic capability by enabling sensitive detection of viral RNA and facilitating molecular epidemiological investigations, including viral lineage identification and transmission tracking (Gigante et al., 2021; Nadin-Davis et al., 2023). In many resource-limited settings, however, traditional histopathological methods such as Seller’s staining continue to be used because they are inexpensive, technically simple, and require minimal laboratory infrastructure (Mananggit et al., 2021; Gavina et al., 2023; Vasconcelos-Martins et al., 2026). Although these conventional methods have lower diagnostic sensitivity than FAT and molecular assays, they remain valuable where access to advanced diagnostic technologies is constrained. The complementary use of conventional and modern diagnostic methods can substantially improve case confirmation and strengthen surveillance systems in endemic regions.

Ghana continues to experience endemic dog-mediated rabies despite decades of control efforts. Human exposures resulting from dog bites are frequently reported across the country, while sporadic outbreaks continue to occur in both urban and rural communities (Bonney et al., 2021). Although national rabies surveillance has improved in recent years, important challenges remain, including inadequate vaccination coverage among dogs, inconsistent reporting of bite incidents, delayed laboratory confirmation, and limited integration between veterinary and public health surveillance systems (Emikpe et al., 2024; Global Alliance for Rabies Control, 2026). These limitations hinder accurate estimation of disease burden and reduce the effectiveness of targeted intervention strategies.

Within the Bono East Region, domestic dogs are commonly kept for security, hunting, and companionship, while free-roaming dog populations remain widespread in many communities. These conditions increase opportunities for virus transmission among animals and elevate the risk of human exposure. Until recently, laboratory confirmation of suspected rabies cases in the region relied largely on Seller’s staining because of limited access to advanced diagnostic technologies. Subsequent investments in diagnostic capacity and collaboration with national reference laboratories have enabled the routine use of FAT and molecular methods, thereby improved the reliability of laboratory confirmation and strengthened regional surveillance activities. Nevertheless, comprehensive analyses describing the epidemiological characteristics of laboratory-confirmed rabies cases in this region remain scarce.

Most published studies from Ghana have focused on human rabies cases, dog bite incidents, or vaccination campaigns. At the same time, relatively few have examined laboratory-confirmed animal cases using routinely collected surveillance data. Information describing temporal trends, vaccination history, animal ownership, species distribution, geographical occurrence, and laboratory confirmation of suspected cases is essential for understanding transmission dynamics and identifying priority areas for intervention. Such evidence is particularly important for evaluating progress towards national rabies elimination goals and informing evidence-based policy within the One Health framework.

The present study addresses this important knowledge gap by analysing six years of laboratory surveillance data from suspected rabies cases submitted to the Regional Veterinary Laboratory in Bono East Region, Ghana. Using a combination of Seller’s staining, FAT, and polymerase chain reaction, we characterised the epidemiological profile of laboratory-confirmed rabies cases between 2020 and 2025. Specifically, the study examined temporal patterns, species distribution, vaccination status, ownership characteristics, bite history, and geographical distribution of submitted animals. By integrating laboratory diagnostic findings with epidemiological surveillance data, this study provides robust evidence of the ongoing burden of canine rabies in Ghana and highlights priorities for strengthening surveillance, improving vaccination coverage, and supporting integrated One Health strategies to achieve the global target of eliminating dog-mediated human rabies by 2030.

### Conceptual One Health Framework for Rabies Transmission

The One Health framework recognizes the interconnected relationships among animal health, human health, and environmental factors in rabies transmission. Integrated approaches combining mass dog vaccination, surveillance, and access to post-exposure prophylaxis are essential for controlling rabies in endemic regions. Studies across Africa have demonstrated that coordinated One Health strategies significantly improve vaccination coverage and disease reporting.

This conceptual model illustrates the interactions among animals, humans, and environmental drivers that influence rabies transmission, as shown in *Figure 1*.

**Figure 1:**
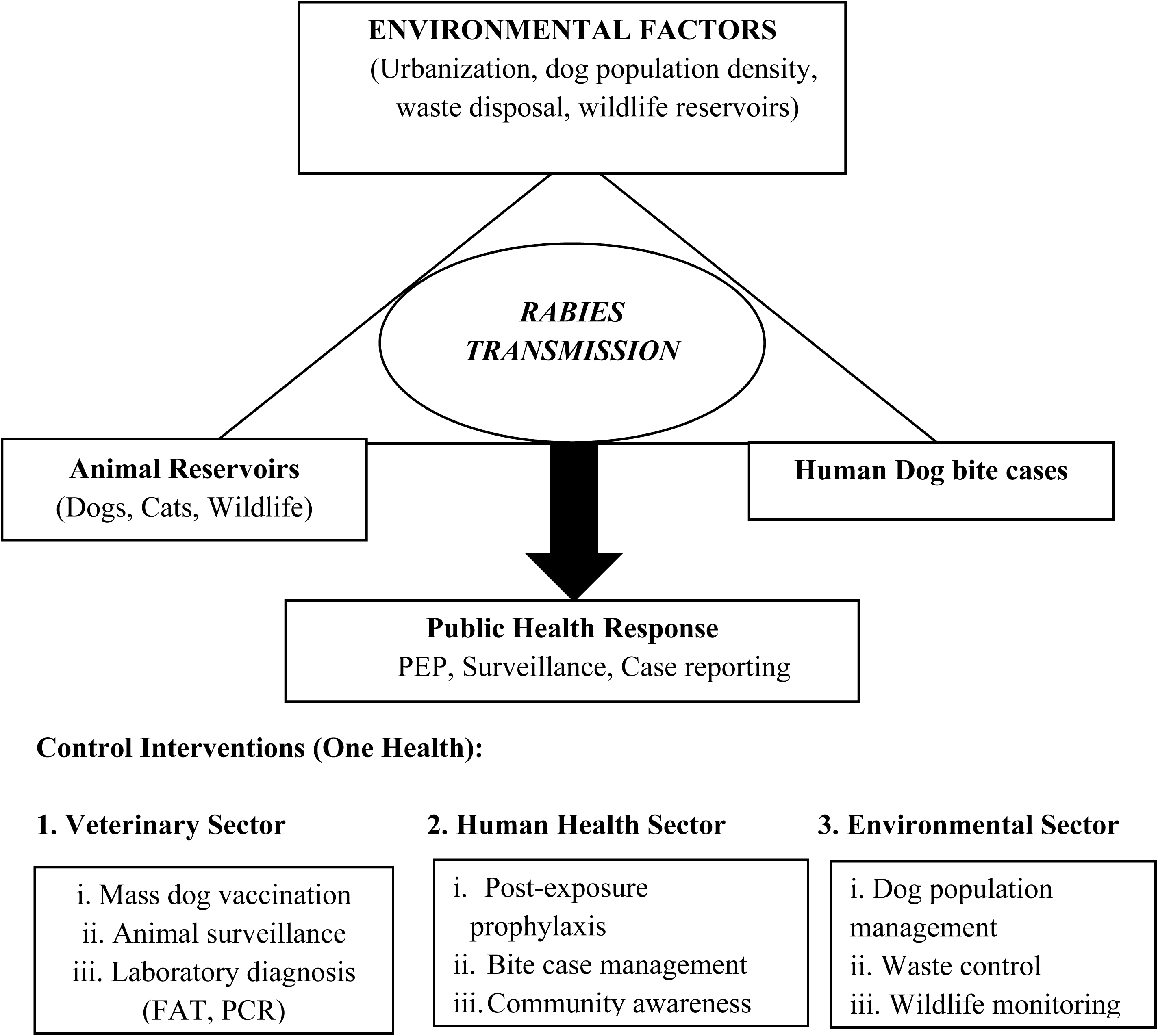
Conceptual One Health Framework Illustrating Transmission Dynamics and Control Pathways of Rabies at the Human-Animal-Environment Interface.

## 2. Materials and Methods

### 2.1. Study Area

This study was conducted at the Regional Veterinary Laboratory, Techiman, which serves the Bono East Region and provides diagnostic support for surrounding districts within the middle belt of Ghana. The region comprises a mixture of urban, peri-urban, and rural communities where livestock production and dog ownership are integral to household livelihoods. Domestic dogs are commonly kept for security, hunting, and companionship, while free-roaming dogs are frequently observed in both urban and rural settlements. The close interaction between dogs, livestock, wildlife, and humans creates favourable conditions for the maintenance and transmission of the rabies virus. The Regional Veterinary Laboratory serves as the primary facility responsible for the investigation and laboratory confirmation of suspected animal rabies cases submitted through the national veterinary surveillance system.

### 2.2. Study Design

A retrospective descriptive study was conducted using routinely collected laboratory surveillance records of animals submitted for rabies diagnosis between January 2020 and December 2025. The study analysed epidemiological and laboratory data from suspected rabies cases received by the Regional Veterinary Laboratory as part of the routine animal disease surveillance programme implemented by the Veterinary Services Directorate. The objective was to describe the epidemiological characteristics of suspected rabies cases and determine the proportion of laboratory-confirmed infections using conventional and reference diagnostic methods.

### 2.3. Case definition and sample collection

A suspected rabies case was defined as any animal presenting with clinical signs compatible with rabies, including aggression, excessive salivation, behavioural changes, abnormal vocalisation, paralysis, incoordination, or other neurological manifestations, and/or an animal with a documented history of biting humans or other animals. This case definition was consistent with recommendations of the WHO (2023) and the WOAH (2023).

Brain tissue specimens were collected only after the death or humane euthanasia of suspected animals by authorised veterinary officers as part of routine disease investigation. Sampling was performed using standard biosafety procedures to minimise occupational exposure to the rabies virus. Brain tissues were obtained from the hippocampus, cerebellum, and brainstem, which are recognised as preferred anatomical sites because of their high viral load in infected animals (WOAH, 2023). Specimens were maintained under an appropriate cold chain during transportation and submitted promptly to the laboratory for diagnostic testing.

#### 2.3.1 Inclusion and exclusion criteria Inclusion criteria

All animal submissions were eligible for inclusion if they met the following criteria:

i. the animal was clinically suspected of rabies based on neurological signs and/or a documented history of biting humans or other animals;
ii. brain tissue specimens or the head or the whole cadaver were submitted to the Regional Veterinary Laboratory between January 2020 and December 2025 for laboratory confirmation;
iii. laboratory records contained sufficient epidemiological information, including species, date of submission, and diagnostic outcome;
iv. the submitted specimen was of adequate quality for at least one diagnostic test.

##### Exclusion criteria

Cases were excluded if laboratory records were incomplete or duplicated, if the specimens were unsuitable for diagnostic examination because of severe decomposition, contamination, or inadequate preservation, or if essential epidemiological or laboratory information required for analysis was unavailable. Records that could not be verified because of missing identifiers or inconsistent documentation were also excluded from the study.

### 2.4 Laboratory diagnosis

Rabies diagnosis was undertaken using one or more laboratory methods, depending on the diagnostic capacity available during the study period. These included the seller’s staining, the FAT, and the polymerase chain reaction (PCR).

#### 2.4.1 Seller’s staining

Seller’s staining was employed for the microscopic detection of Negri bodies in brain tissue smears. Impression smears were prepared on clean microscope slides, air-dried, fixed in absolute methanol, stained using Seller’s stain, and examined under a light microscope. The presence of characteristic intracytoplasmic eosinophilic inclusion bodies within neuronal cells was considered suggestive of rabies infection. Although Seller’s staining has historically been widely used in rabies diagnosis, its diagnostic sensitivity is lower than that of antigen- and molecular-based methods (Diallo et al., 2019; Amicizia et al., 2024; Fotedar & Ravish, 2025; Ritchie et al., 2025; Fotedar & Shankariah, 2026; Pardeshi et al., 2026).

#### 2.4.2 FAT

The FAT was used for confirmatory detection of rabies virus antigen in brain tissue samples. Thin brain impression smears were prepared on microscope slides, fixed in chilled acetone, and incubated with fluorescein isothiocyanate-labelled anti-rabies antibodies according to established laboratory protocols. After washing, the slides were examined using a fluorescence microscope. Samples demonstrating characteristic apple-green fluorescence within infected neuronal tissue were interpreted as positive for rabies virus antigen. The FAT remains the internationally recommended reference test for routine post-mortem diagnosis because of its high diagnostic sensitivity and specificity (WHO, 2023; WOAH, 2023).

#### 2.4.3 Polymerase chain reaction (PCR)

Selected brain tissue samples underwent molecular confirmation by reverse transcription polymerase chain reaction (RT-PCR). Viral RNA was extracted from brain tissue using a commercially available RNA extraction kit according to the manufacturer’s instructions. Complementary DNA (cDNA) was synthesised by reverse transcription and subsequently amplified using rabies virus-specific primers. Amplified products were analysed by agarose gel electrophoresis to confirm the presence of rabies virus nucleic acid. Molecular diagnosis provided an additional level of confirmation and enhanced the reliability of laboratory surveillance, particularly in samples with equivocal or inconclusive results (Gigante et al., 2021).

#### 2.4.4 Quality assurance

Quality assurance procedures were implemented throughout specimen collection, transportation, laboratory testing, and data recording to ensure the reliability of diagnostic results. Brain tissue samples were collected by trained veterinary personnel using standard biosafety procedures and transported under a maintained cold chain to preserve sample integrity. Laboratory analyses were performed by qualified personnel following established standard operating procedures for rabies diagnosis. Appropriate positive and negative controls were included during immunofluorescence and molecular assays where applicable, and laboratory findings were independently reviewed before final reporting. Diagnostic procedures were conducted in accordance with internationally recognised recommendations for rabies diagnosis published by the WOAH (2023) and the WHO (2023).

#### 2.4.5. Capacity strengthening and laboratory collaboration

During the early phase of the study (2020 – 2022), routine laboratory diagnosis of suspected rabies cases at the Regional Veterinary Laboratory relied primarily on Seller’s staining because of limited access to advanced diagnostic infrastructure. In 2022, laboratory personnel and regional veterinary officers participated in a national rabies capacity-building workshop that strengthened technical competencies in rabies surveillance, sample handling, and laboratory diagnosis.

Following this training, the laboratory established collaborations with national reference laboratories to improve diagnostic confirmation of suspected rabies cases. Initially, brain tissue specimens were referred to the Kumasi Centre for Collaborative Research in Tropical Medicine (KCCR), Kumasi, for confirmatory testing using internationally recommended diagnostic methods, which include molecular assays. Subsequently, from 2024 onward, samples were then referred to either the Accra Veterinary Laboratory, the Kumasi Veterinary Laboratory, or the Central Veterinary Laboratory for confirmatory diagnosis. These collaborations substantially improved access to quality-assured laboratory services, strengthened regional rabies surveillance, enhanced diagnostic accuracy, and supported timely reporting of confirmed cases.

### 2.5. Data Collection

Laboratory records were reviewed to extract relevant epidemiological and diagnostic information for each submitted case. The variables collected included animal species, vaccination status, ownership status (owned or stray), history of biting humans or other animals, laboratory diagnostic results and date of sampling, submission and diagnosis The extracted data were compiled into a structured dataset for further analysis.

### 2.6. Data Analysis

Data were analysed using descriptive statistical methods. Categorical variables were summarised as frequencies and percentages, whereas continuous variables were summarised using appropriate descriptive measures where applicable. The laboratory-confirmed positivity rate was calculated as the proportion of confirmed rabies cases among all animals submitted for diagnostic investigation during the study period.

Data management and statistical summaries were performed using Microsoft Excel 2019 (Microsoft Corporation, Redmond, WA, USA). Tables and graphical presentations were generated to illustrate temporal trends and the distribution of epidemiological variables. Because the study was descriptive and based on routine surveillance data, no inferential statistical analyses were undertaken.

### 2.7. Ethical Considerations

The study analysed anonymised secondary data generated through the routine rabies surveillance programme of the Veterinary Services Directorate. No animals were handled or euthanised specifically for research purposes, and all specimens originated from routine diagnostic investigations undertaken by authorised veterinary personnel. Confidentiality of surveillance records was maintained throughout data extraction, analysis, and reporting.

Administrative permission to access and analyse the laboratory surveillance records was obtained from the Veterinary Services Directorate, Ministry of Food and Agriculture, Ghana (*Administrative Permission Reference No.:* BE/RVO/RABV-RES09).

The study complied with national guidelines governing the use of surveillance data for research and was conducted in accordance with internationally accepted ethical principles for veterinary epidemiological investigations.

### 2.9 Reporting guideline

The study was designed, analysed, and reported in accordance with the Strengthening the Reporting of Observational Studies in Epidemiology, Veterinary Extension (STROBE-Vet) statement, which provides recommendations for transparent reporting of observational studies involving animal populations (Sargeant et al., 2016). The STROBE-Vet recommendations were considered throughout study design, data analysis, and manuscript preparation to improve methodological transparency, reproducibility, and completeness of reporting.

## 3. Result

### 3.1 Characteristics of submitted rabies cases

Between January 2020 and December 2025, the Regional Veterinary Laboratory received 51 brain tissue specimens from animals suspected of rabies for laboratory confirmation. The annual number of submissions ranged from one case in 2023 to 14 cases in 2025 (Table 1). Of the 51 submitted specimens, 46 were laboratory-confirmed as rabies, yielding an overall positivity rate of 90.2% (95% confidence interval [CI]: 78.6% – 96.7%). Five samples (9.8%) tested negative.

**Table 1:**
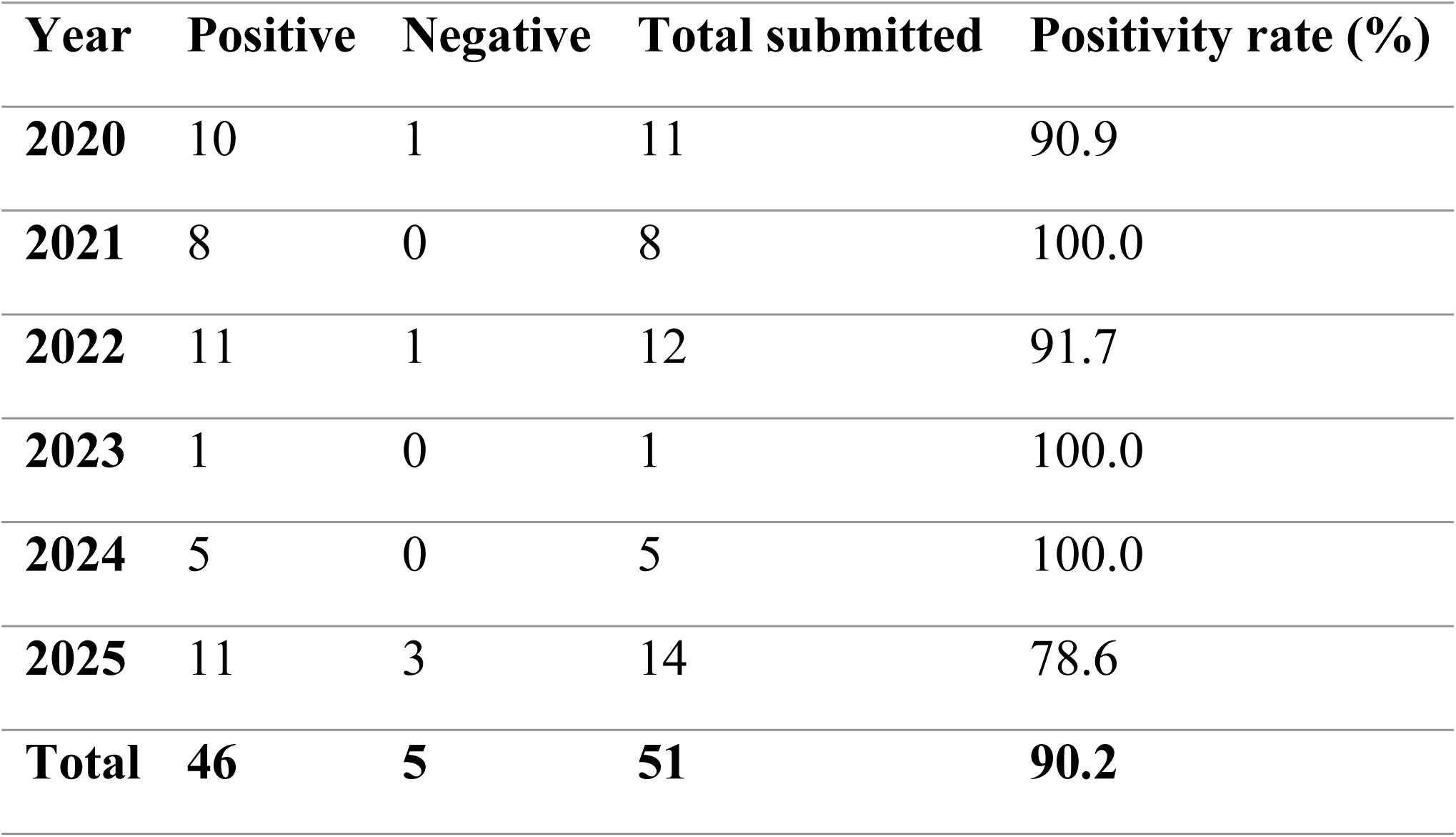
Annual laboratory investigation of suspected rabies cases, 2020 – 2025.

| Year | Positive | Negative | Total submitted | Positivity rate (%) |
| --- | --- | --- | --- | --- |
| 2020 | 10 | 1 | 11 | 90.9 |
| 2021 | 8 | 0 | 8 | 100.0 |
| 2022 | 11 | 1 | 12 | 91.7 |
| 2023 | 1 | 0 | 1 | 100.0 |
| 2024 | 5 | 0 | 5 | 100.0 |
| 2025 | 11 | 3 | 14 | 78.6 |
| Total | 46 | 5 | 51 | 90.2 |

### 3.2 Laboratory Diagnosis of Suspected Rabies Cases

A total of 51 suspected rabies cases were submitted to the veterinary diagnostic laboratory for confirmation during the study period. Brain tissue samples from these animals were analyzed using Seller’s staining technique, FAT, and PCR for the detection of the Rabies virus. Out of the 51 samples examined, 90.20% (46/51) were laboratory-confirmed positive, while the remaining 9.80% (5/51) tested negative, as shown in *Figure 2*.

**Figure 2:**
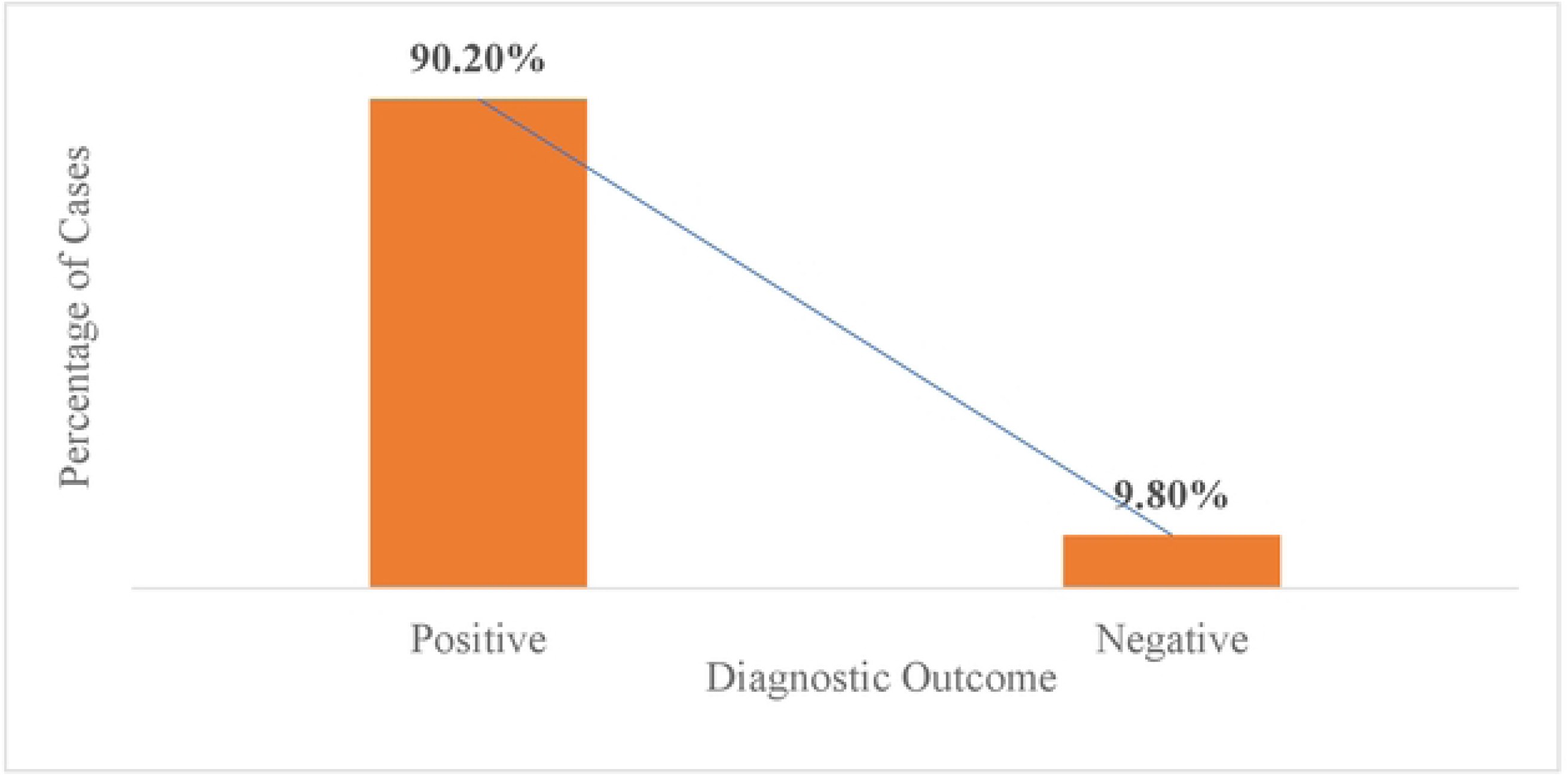
Laboratory Diagnosis of Suspected Rabies Cases.

The diagnostic approach evolved during the surveillance period. Prior to 2022, laboratory confirmation relied primarily on the Seller’s staining. Following the implementation of enhanced diagnostic capacity, suspected cases were increasingly confirmed using FAT and RT-PCR. As shown in Table 2, in 2022 all 12 samples were tested with Seller’s staining whiles 10 and 5 were tested with RT-PCR and FAT, respectively.

**Table 2:** Diagnostic methods used during surveillance.

| Year | Seller's stain | FAT | RT-PCR |
| --- | --- | --- | --- |
| 2020 | 10 | 0 | 0 |
| 2021 | 8 | 0 | 0 |
| 2022 | 12 | 5 | 10 |
| 2023 | 0 | 1 | 1 |
| 2024 | 0 | 5 | 5 |
| 2025 | 0 | 14 | 14 |

#### Distribution of Suspected Rabies Cases by Animal Species

Analysis of the species distribution revealed that domestic dogs accounted for the majority of submissions. From *Figure 3,* out of the total samples analyzed, 49 cases (96.10%) were from dogs, while 2 cases (3.90%) involved cats.

**Figure 3:**
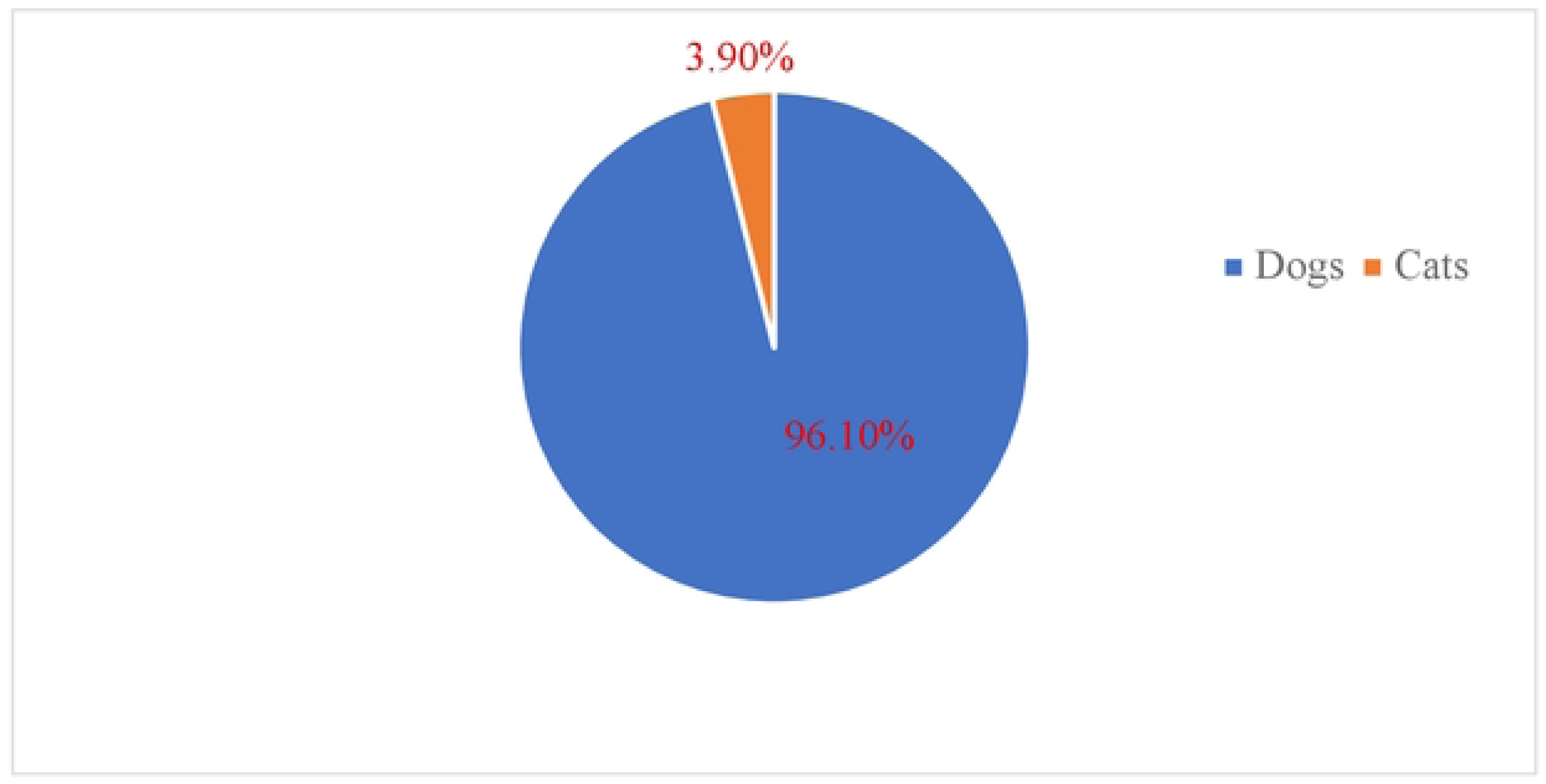
Animal Species Distribution.

### 3.4 Vaccination Status of Suspected Animals

Vaccination history was assessed to evaluate the potential role of vaccination gaps in rabies transmission, as shown in *Figure 4*. The results revealed that 64.70% (33) of the animals were reported as unvaccinated, while33.30% (17/51) had an unknown vaccination status. Only one animal (2.00%) was reported to have been vaccinated against rabies.

**Figure 4:**
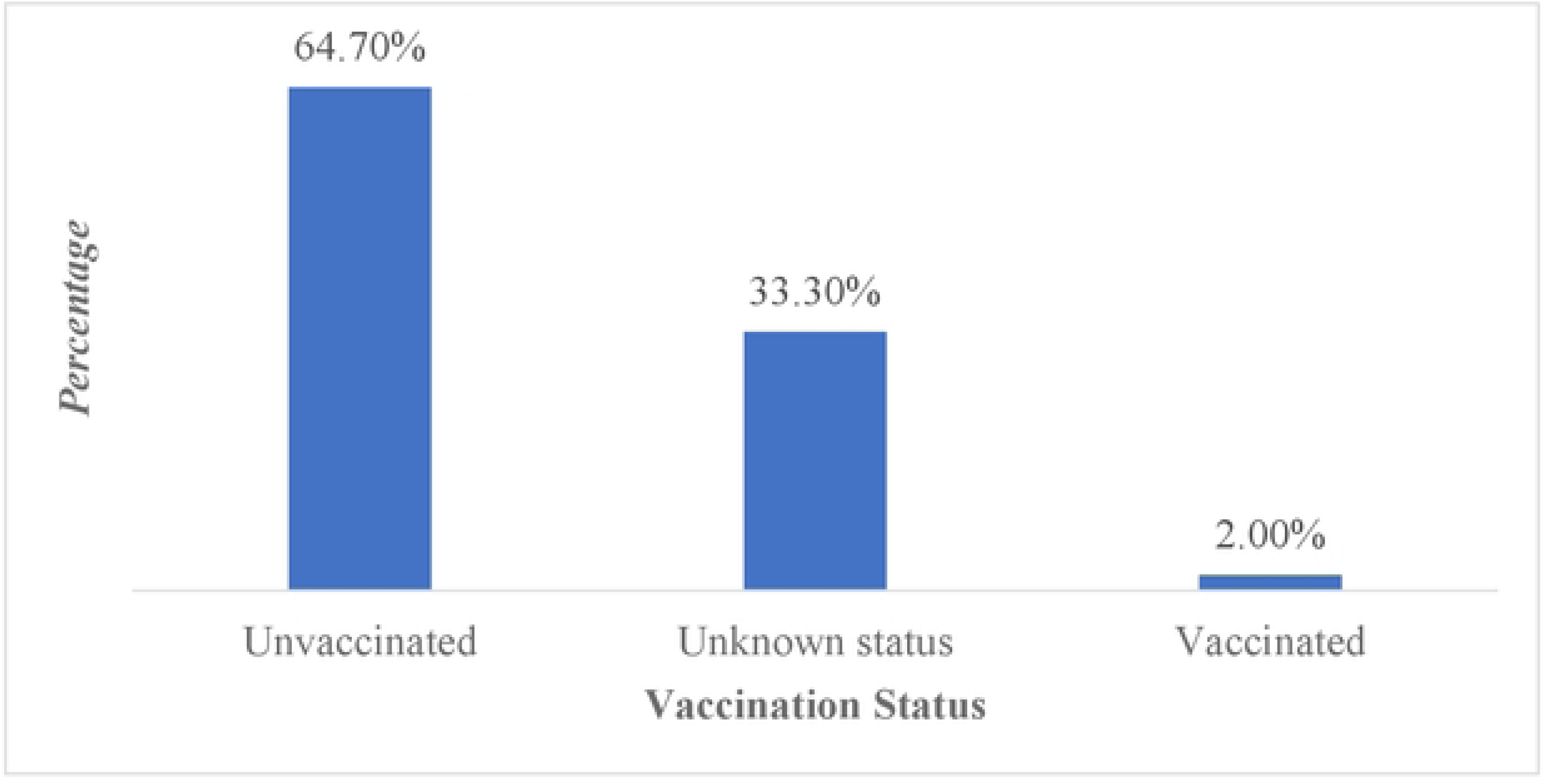
Vaccination Status of Suspected Animals.

### 3.5 Human Bite Exposure Associated with Suspected Rabies Cases

A large proportion of animals submitted for rabies testing had a documented history of biting humans. 46 animals (90.2%) were reported to have bitten one or more people before submission for laboratory testing, while 5 animals (9.8%) had no reported history of biting humans but had been bitten by other animals or had bitten other animals, as shown in Table 3.

**Table 3:** Bite Exposure History.

| Bite History | Number | Percentage (%) |
| --- | --- | --- |
| Bit humans/animal | 46 | 90.2 |
| No bite reported | 5 | 9.8 |

### 3.6 Geographic distribution

Analysis of the geographic origin of the submitted samples revealed that most suspected rabies cases originated from communities in the Techiman area. Additional cases were reported from surrounding districts within the region. *Figure 5* illustrates the geographic locations of animals submitted for laboratory diagnosis of rabies. Most cases were clustered within communities surrounding Techiman, including Jamestown-Techiman, Kenten-Techiman, Aworano-Techiman, and other peri-urban settlements. Additional cases were reported from surrounding towns, including Sunyani, Sunyani-Odumasi, Dormaa Ahenkro, Kintampo, Yeji, and Bechem. The spatial clustering of suspected cases around Techiman suggests localized transmission dynamics and highlights priority areas for targeted rabies surveillance and mass dog vaccination campaigns.

**Figure 5:**
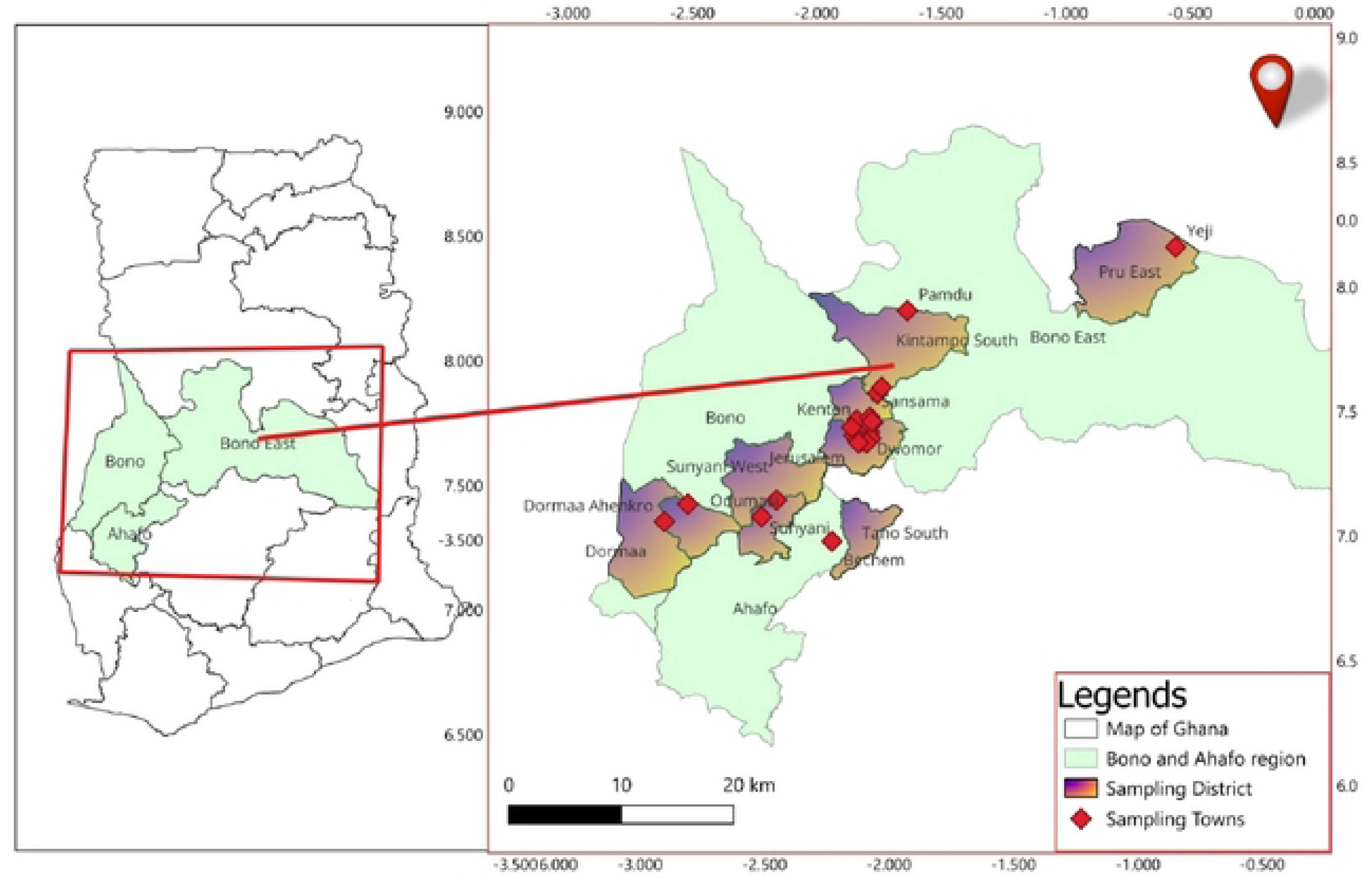
Spatial distribution of suspected rabies cases in the Study area.

#### 3.6.1 District distribution of laboratory-confirmed rabies cases

Laboratory-confirmed rabies cases were reported from multiple districts within the laboratory catchment area. Table 4 showed that Techiman North and Techiman South contributed the largest number of submissions, 19.8 and 62.8, respectively, while fewer cases originated from more distant districts. Laboratory-confirmed cases were identified across several administrative areas, demonstrating widespread occurrence of suspected rabies within the surveillance network.

**Table 4:** Distribution of laboratory-confirmed rabies cases by location (2020 – 2025)

| District | Suspected | Confirmed | Positivity (%) |
| --- | --- | --- | --- |
| Techiman North | 10 | 10 | 19.9 |
| Techiman South | 32 | 32 | 62.8 |
| Kintampo South | 1 | 1 | 1.9 |
| Sunyani East | 1 | 1 | 1.9 |
| Sunyani West | 2 | 2 | 3.9 |
| Dormaa Central | 2 | 2 | 3.9 |
| Dormaa East | 1 | 1 | 1.9 |
| Tano South | 1 | 1 | 1.9 |
| Pru East | 1 | 1 | 1.9 |

### 3.7 Association between epidemiological characteristics and laboratory-confirmed rabies

Domestic dogs accounted for 49 (96.1%) of the submitted animals, of which 44 (89.8%) were laboratory confirmed, whereas both cats submitted during the study period tested positive. Among unvaccinated animals, 28 of 33 (84.8%) were laboratory confirmed, while all animals with documented vaccination (1/1) or unknown vaccination status (17/17) tested positive. A history of biting humans was recorded for 46 animals, of which 42 (91.3%) were laboratory confirmed from Table 5. Although descriptive differences were observed across categories, the small number of negative cases limited the statistical power to detect meaningful associations. Consequently, Fisher’s exact test was used to evaluate these relationships, and the findings should be interpreted cautiously. The Pearson chi-square test is presented because it is widely available in standard statistical software.

**Table 5:** Association between selected epidemiological characteristics and laboratory-confirmed rabies among suspected animal cases submitted to the Regional Veterinary Laboratory, 2020–2025.

| Variable | Category | Rabies positive n (%) | Rabies negative n (%) | Total (n) | Statistical test | P-value |
| --- | --- | --- | --- | --- | --- | --- |
| Species | Dog | 44 (89.8) | 5 (10.2) | 49 | Fisher's exact test | 1.000 |
|  | Cat | 2 (100.0) | 0 (0.0) | 2 |  |  |
| Vaccination status | Vaccinated | 1 (100.0) | 0 (0.0) | 1 | Pearson's $\chi^2$ test† | 0.220 |
|  | Unvaccinated | 28 (84.8) | 5 (15.2) | 33 |  |  |
|  | Unknown | 17 (100.0) | 0 (0.0) | 17 |  |  |
| History of human bite | Yes | 42 (91.3) | 4 (8.7) | 46 | Fisher's exact test | 0.416 |
|  | No | 4 (80.0) | 1 (20.0) | 5 |  |  |

**Abbreviations:** $\chi^2$ , chi-square; $n$ , number of animals. † Owing to sparse cell counts, an exact Fisher-Freeman-Halton test is preferable where available. Fisher's exact test was used where expected cell frequencies were <5.

### 3.8 Summary of Key Epidemiological Findings

Overall, the laboratory surveillance data demonstrate a high positivity rate among suspected rabies cases, with domestic dogs representing the predominant species involved. The majority of suspected animals were either unvaccinated or had unknown vaccination status, and a substantial proportion had a history of biting humans, underscoring the ongoing zoonotic risk posed by rabies within the study area.

These findings provide valuable epidemiological evidence to support improved rabies control strategies, particularly mass dog vaccination campaigns, strengthened surveillance systems, and enhanced public awareness of rabies prevention.

## Discussion

This study provides recent laboratory-based epidemiological evidence on animal rabies in the Bono East Region of Ghana and offers important evidence on the persistence of canine rabies within the country’s middle belt. Using six years of routine surveillance data, the study demonstrated a consistently high laboratory confirmation rate among suspected rabies cases, with domestic dogs accounting for nearly all confirmed infections. Furthermore, most affected animals were either unvaccinated or had no documented vaccination history, and the majority had reportedly bitten humans before laboratory investigation. Collectively, these findings reinforce the continued importance of domestic dogs as the principal reservoir and source of human rabies exposure in Ghana and support ongoing national and global efforts aimed at eliminating dog-mediated human rabies.

One of the most striking findings of this study was the overall laboratory positivity rate of 90.2% among suspected rabies cases. This exceptionally high confirmation rate suggests that veterinary officers were generally able to identify clinically compatible cases before laboratory submission. Similar high positivity rates have been reported in surveillance studies from several African countries where samples submitted for diagnosis usually originate from animals exhibiting advanced neurological signs or those implicated in human bite incidents (Coetzer et al., 2019; Sambo et al., 2022; Keshavamurthy et al., 2024; Kahariri et al., 2025). The high confirmation rate observed in the present study also reflects improvements in diagnostic capacity following the introduction of reference laboratory support using the FAT and RT-PCR, both of which have superior diagnostic performance compared with conventional microscopic techniques (Diallo et al., 2019; Nadin-Davis et al., 2023; Amicizia et al., 2024; Fotedar & Ravish, 2025; Ritchie et al., 2025; Fotedar & Shankariah, 2026; Pardeshi et al., 2026).

Rabies cases were detected throughout every year of the surveillance period, indicating persistent viral circulation within the study area. Although annual case numbers fluctuated, confirmed infections were recorded continuously between 2020 and 2025, demonstrating that rabies remains endemic within the laboratory catchment area. Such year-to-year variation is expected in passive surveillance systems because submission rates are influenced by public awareness, access to veterinary services, reporting practices, and laboratory capacity rather than true disease incidence alone. Similar temporal fluctuations have been documented in Ghana and elsewhere in sub-Saharan Africa, where under-reporting and inconsistent surveillance remain important obstacles to accurately estimating rabies burden (Bonney et al., 2021; Taylor et al., 2022; WHO, 2023).

The marked decline in confirmed cases observed during 2023 should therefore be interpreted cautiously. Rather than indicating a genuine reduction in rabies transmission, the low number of laboratory-confirmed cases may reflect reduced sample submission, changes in surveillance intensity, logistical challenges affecting specimen transport, or variations in community reporting behaviour. Passive surveillance systems frequently underestimate the true burden of rabies because many suspect animals are never submitted for laboratory confirmation, particularly in rural and resource-limited settings (Coetzer et al., 2019; Nel et al., 2021; Keshavamurthy et al., 2024; Kahariri et al., 2025). Consequently, laboratory-confirmed cases likely represent only a fraction of infections occurring within the wider population.

The study further demonstrated the overwhelming predominance of domestic dogs, which accounted for 96.1% of all submitted animals. This finding is consistent with the established epidemiology of rabies in Africa, where domestic dogs remain responsible for more than 95% of human rabies exposures and serve as the principal maintenance host for rabies virus transmission (Hampson et al., 2022; WHO, 2023). The limited number of feline submissions in this study most likely reflects spill-over infection from infected dogs rather than independent maintenance of rabies virus within cat populations. Similar observations have been reported in studies from other endemic countries, where infections in cats occur sporadically following exposure to rabid dogs and contribute minimally to sustained transmission (Wallace et al., 2022).

The continued dominance of canine rabies observed in the present study has important implications for disease control. Mathematical modelling and field evidence consistently demonstrate that sustained vaccination of at least 70% of the dog population is sufficient to interrupt rabies transmission under most epidemiological conditions (Lechenne et al., 2021; Undurraga et al., 2023). Consequently, effective control programmes should prioritize comprehensive mass dog vaccination campaigns, supported by responsible dog ownership, improved registration systems, and strengthened veterinary surveillance. Such interventions remain considerably more cost-effective than relying primarily on human post-exposure prophylaxis after exposure has occurred (Undurraga et al., 2023).

The transition in diagnostic practice observed during the study period represents another important strength of the regional surveillance system. Before 2022, diagnosis depended largely on Seller’s staining because of limited laboratory infrastructure. Although historically valuable, Seller’s staining has substantially lower sensitivity than antigen- and molecular-based diagnostic methods and may fail to detect infected animals with low viral antigen concentrations (Diallo et al., 2019; WOAH, 2023; Amicizia et al., 2024; Fotedar & Ravish, 2025; Ritchie et al., 2025; Fotedar & Shankariah, 2026; Pardeshi et al., 2026). Following national capacity-building initiatives, suspected rabies samples were increasingly referred to accredited reference laboratories for confirmation using FAT and RT-PCR. This transition likely enhanced diagnostic accuracy, reduced the risk of false-negative diagnoses, and improved confidence in surveillance data. The successful collaboration between the Regional Veterinary Laboratory, the KCCR, and national veterinary reference laboratories illustrates the value of inter-institutional partnerships in strengthening rabies surveillance in resource-constrained settings.

The high diagnostic confirmation rate achieved through this collaborative network also demonstrates the importance of sustained investment in laboratory infrastructure and workforce development. Reliable laboratory confirmation not only improves surveillance quality but also informs timely public health responses, including contact tracing, risk assessment, and decisions regarding administration of post-exposure prophylaxis following human exposure. As countries progress toward the global goal of eliminating dog-mediated human rabies by 2030, strengthening diagnostic capacity will remain an essential component of effective national rabies control programmes (FAO, WHO & WOAH, 2024).

An important finding of this study was the low documented rabies vaccination coverage among animals submitted for laboratory diagnosis. Nearly two-thirds of the animals had no history of rabies vaccination, while the vaccination status of approximately one-third could not be verified. Only one animal was reported to have received rabies vaccination before disease onset. Although statistical analysis did not demonstrate a significant association between vaccination status and laboratory-confirmed rabies, this finding should be interpreted with caution because the study included only one vaccinated animal and a limited number of laboratory-negative cases, thereby reducing the statistical power to detect meaningful differences. Nevertheless, the predominance of unvaccinated animals among confirmed cases is consistent with previous reports from Ghana and other rabies-endemic countries, where inadequate vaccination coverage remains a major driver of sustained virus transmission (Bonney et al., 2021; WHO, 2023).

The substantial proportion of animals with unknown vaccination histories also highlights weaknesses in animal identification and vaccination record-keeping. In many low- and middle-income countries, routine documentation of canine vaccination remains incomplete, particularly among free-roaming and community-owned dogs. This limitation complicates surveillance activities and reduces the ability of veterinary authorities to evaluate the effectiveness of vaccination programmes. Strengthening dog registration systems, improving vaccination certification, and implementing electronic surveillance databases would facilitate more accurate monitoring of vaccination coverage and support evidence-based planning of rabies control interventions (Banyard et al., 2022; FAO, WHO & WOAH, 2024).

Human exposure represented another prominent epidemiological feature of the present study. More than 90% of submitted animals had reportedly bitten one or more persons before laboratory investigation, indicating that most laboratory submissions were initiated following potential human exposure. This finding underscores the continuing public health importance of canine rabies in the Bono East Region and demonstrates the close epidemiological relationship between animal rabies surveillance and human rabies prevention. Similar observations have been reported throughout Africa, where human exposure frequently serves as the principal trigger for laboratory investigation of suspected rabid animals (Taylor et al., 2022; Sambo et al., 2022). These findings further emphasise the importance of prompt laboratory confirmation in guiding risk assessment and supporting timely administration of post-exposure prophylaxis to exposed individuals.

The geographical distribution of laboratory-confirmed rabies cases demonstrated that infections were reported across several districts within the laboratory catchment area, although nearly half of all confirmed cases originated from the Techiman Municipality. This concentration may reflect the relatively high human and dog population density within Techiman, greater accessibility to veterinary services, and the location of the Regional Veterinary Laboratory, which likely facilitates specimen submission from nearby communities. Conversely, the comparatively small number of submissions from more distant districts should not necessarily be interpreted as indicating lower disease occurrence. Limited access to diagnostic facilities, transportation challenges, reduced public awareness, and under-reporting may contribute to lower submission rates from peripheral communities, a pattern that has been widely documented in passive rabies surveillance systems across sub-Saharan Africa (Coetzer et al., 2019; Nel et al., 2021; Keshavamurthy et al., 2024; Kahariri et al., 2025).

The statistical analyses performed in this study did not identify significant associations between laboratory-confirmed rabies and animal species, vaccination status, or reported history of human bite. However, these findings should not be interpreted as evidence that these factors are unrelated to rabies occurrence. Instead, the absence of statistical significance most likely reflects the limited sample size, the overwhelming predominance of positive cases, and sparse observations within several comparison groups. Small datasets frequently have insufficient statistical power to detect modest epidemiological associations, particularly when one outcome category contains relatively few observations. Future investigations involving larger multicentre datasets would permit more robust analytical approaches, including multivariable regression models capable of identifying independent risk factors for laboratory-confirmed rabies.

The strengthened laboratory network established during the study period represents a notable achievement for rabies surveillance within the Bono East Region. The referral of specimens to accredited reference laboratories for confirmation by FAT and RT-PCR substantially improved diagnostic confidence and aligned regional surveillance practices with internationally recommended standards (WOAH, 2023; WHO, 2023). Such collaborations illustrate how strategic investment in laboratory infrastructure, workforce development, and institutional partnerships can strengthen surveillance capacity in resource-limited settings. Continued support for these collaborative networks will be essential for achieving reliable disease detection and monitoring progress towards national rabies elimination targets.

The findings of this study reinforce the importance of adopting a One Health approach to rabies prevention and control. Rabies remains a disease that exists at the interface of human, animal, and environmental health, and its effective control depends upon coordinated collaboration among veterinary services, public health authorities, local governments, wildlife agencies, and affected communities (Cleaveland et al., 2021; Destoumieux-Garzón et al., 2022). Laboratory surveillance should therefore be integrated with community-based education, responsible dog ownership initiatives, mass canine vaccination campaigns, improved access to human post-exposure prophylaxis, and strengthened reporting systems. Such integrated approaches have consistently been shown to reduce rabies transmission and represent the cornerstone of the global strategy to eliminate dog-mediated human rabies by 2030 (FAO, WHO & WOAH, 2024; Lechenne et al., 2021).

This study has several strengths. It provides six years of laboratory-confirmed surveillance data from a region where published epidemiological information on animal rabies remains limited. The use of internationally recognised diagnostic methods, including FAT and RT-PCR, enhances the reliability of the reported findings. Furthermore, the study documents the successful transition from conventional microscopy to modern reference laboratory diagnosis, illustrating the impact of capacity strengthening on regional surveillance systems. The inclusion of geographical information and epidemiological characteristics also provides valuable baseline data for future rabies control programmes within Ghana.

Several limitations should also be acknowledged. First, the retrospective design relied on routinely collected surveillance records, which limited the availability of detailed demographic and exposure information for some animals. Second, passive surveillance is inherently susceptible to under-reporting because only animals submitted for laboratory investigation were included, and many suspected cases in remote communities may never have been sampled. Third, the relatively small sample size restricted the statistical power of analytical comparisons and limited the ability to identify independent risk factors associated with laboratory-confirmed rabies. Finally, because the study was laboratory-based, the findings cannot be interpreted as estimates of the true incidence or prevalence of rabies within the wider animal population.

Despite these limitations, the study provides important epidemiological evidence demonstrating the continued circulation of the rabies virus within the Bono East Region and highlights critical gaps in canine vaccination coverage and surveillance. The predominance of domestic dogs among laboratory-confirmed cases, together with the high frequency of human exposure events, emphasises the continuing public health importance of canine rabies in Ghana. Sustained investment in mass dog vaccination, integrated One Health surveillance, laboratory diagnostic capacity, community engagement, and equitable access to post-exposure prophylaxis will be essential for accelerating progress toward the elimination of dog-mediated human rabies in Ghana and achieving the global Zero by 30 targets.

## Acknowledgments

The authors sincerely acknowledge the technical support provided by the Kumasi Centre for Collaborative Research in Tropical Medicine for its contribution to the laboratory diagnosis and analysis of suspected Rabies samples used in this study. We also express our appreciation to the staff of the Accra Veterinary Laboratory, Kumasi Veterinary Laboratory, and Central Veterinary Laboratory-Pong Tamale for their assistance in receiving, processing, and analysing the samples through established diagnostic techniques, including fluorescent antibody testing and polymerase chain reaction.

The authors are further grateful to the field and laboratory personnel of the Disease Investigation Farm and Veterinary Laboratory under the Veterinary Services Directorate in Ghana for their efforts in surveillance activities, sample collection, and data management. Special appreciation is extended to all officers and collaborators who supported rabies surveillance and reporting within the Bono East Region, which made this study possible.

## Author Contributions

**PKD:** Conceptualization, Methodology, Formal analysis, Investigation, Data curation, Writing-original draft preparation

**SAS:** Conceptualization, Methodology, Validation, Writing – review & editing

**AO-T, CANN & PA:** Formal analysis, Investigation, Writing – review & editing

**RTG, SO-W & EO:** Investigation, Supervision, Software, Writing – review & editing

**IB**: Data curation, Project administration, Resources, Writing – review & editing

**HD-F, AN, DK & NYA-B:** Supervision, Validation, Project administration, Writing – review & editing

All authors have read and agreed to the published version of the manuscript.

## Informed Consent Statement

Not applicable.

## Data Availability Statement

All relevant data supporting the findings of this study are publicly available at DOI: 10.5281/zenodo.21212788.

## Abbreviations

cDNA: Complementary Deoxyribonucleic Acid
CI: Confidence Interval
FAO: Food and Agriculture Organization of the United Nations
FAT: Fluorescent Antibody Test
KCCR: Kumasi Centre for Collaborative Research in Tropical Medicine
PCR: Polymerase Chain Reaction
PEP: Post-Exposure Prophylaxis
RABV: Rabies Virus
RNA: Ribonucleic Acid
RT-PCR: Reverse Transcription Polymerase Chain Reaction
STROBE-Vet: Strengthening the Reporting of Observational Studies in Epidemiology–Veterinary Extension
VSD: Veterinary Services Directorate
WHO: World Health Organization
WOAH: World Organisation for Animal Health
χ²: Chi-square test

